# System-wide profiling by proteome integral solubility alteration assay of drug residence times for target characterization

**DOI:** 10.1101/2022.06.27.497697

**Authors:** Pierre Sabatier, Christian M. Beusch, Zhaowei Meng, Roman A. Zubarev

## Abstract

Most drugs used in the clinic and drug candidates target multiple proteins, and thus detailed characterization of their efficacy targets is required. While current methods rely on quantitative measurements at thermodynamic equilibrium, kinetic parameters such as the residence time of a drug on its target provide a better proxy for efficacy *in vivo*. Here, we present Residence Time Proteome Integral Solubility Alteration (ResT-PISA) assay which provides monitoring temporal protein solubility profiles after drug removal (“off-curve”) in cell lysate or intact cells, quantifying the lifetime of drug-target interaction. A compressed version of the assay measures the integral under the off-curve enabling the multiplexing of binding affinity and residence time assessments into a single proteomic analysis. We introduce a combined scoring system for three parametric dimensions to improve prioritization of targets. By providing complementary information to other characteristics of drug-target interaction, ResT-PISA approach will be useful in drug development and precision medicine.

## Introduction

The major disconnect existing between *in vitro* data on drug-target interactions and drug efficacy in humans is one of the primary sources of attrition during drug discovery^1,2^. Traditionally, equilibrium thermodynamic constants measured *in vitro*, such as Kd and Ki, have been used to assess the chances of a drug candidate for efficient target engagement *in vivo*. However, due to the biological complexity of human organism it is unlikely that a single *in vitro* parameter can reliably predict *in vivo* drug efficacy; a model taking into account a number of orthogonal parameters will have an *a priori* higher predictive power.

One of the critical contributors to the drug action mechanism *in vivo* is the lifetime of the drug-target complex, during which the drug molecule affects the biological system. Thus, residence time of the drug on target, which is the reciprocal of the rate constant for dissociation of the drug-target complex, is emerging as a strong contributor to the desired predicting model of *in vivo* efficacy^3–9^. The importance of such a kinetic parameter is due to the fact that cells, organs and organisms are open systems operating far from equilibrium. In open systems the concentrations of the drug, the endogenous ligands for the target and the target itself fluctuate with time^10^. Thus, residence time may correlate with *in vivo* drug efficacy better than equilibrium *in vitro* constants. Indeed, empirical observations support the notion that compounds with longer residence time have higher *in vivo* efficacy^11^.

Another common complicating factor in drug development is that the drug usually interacts *in vivo* with several proteins, the main target as well as off-targets, with all these molecules competing with each other for the ligand in an open system. As a drug only exerts its effect through a given target when it is bound to that target, in order to predict the drug efficacy in such a complex environment one has to be able to compare the lifetimes of all individual drug-target complexes. For example, the difference in residence time of a drug on its target and on the off-target proteins responsible for toxic side-effects defines to large extent the therapeutic window of the drug^10,12,13^.

It has thus been suggested that measurements of drug-target residence time should be incorporated into the standard drug discovery procedure during the lead optimization phase^14^. Such measurements are usually performed either with surface plasmon resonance, where a model fitted to sensogram data provides the residence time value^15^, or with competitive binding assay using designed competitive probes^16^. In the first case the model assumes that drug binding to the pocket is either a simple one-step process (Fig. 1a) or a more complicated two-step process. In the latter case, the initial, weaker binding at or near the pocket is followed by binding-induced structural changes in the enzyme to better accommodate the drug. This accommodation results in stronger binding and consequently longer lifetime of the drug-target complex^17^. The corresponding “off” curves are thus either exponential or exhibit both fast and slow components (Fig. 1b).

**Fig. 1.**
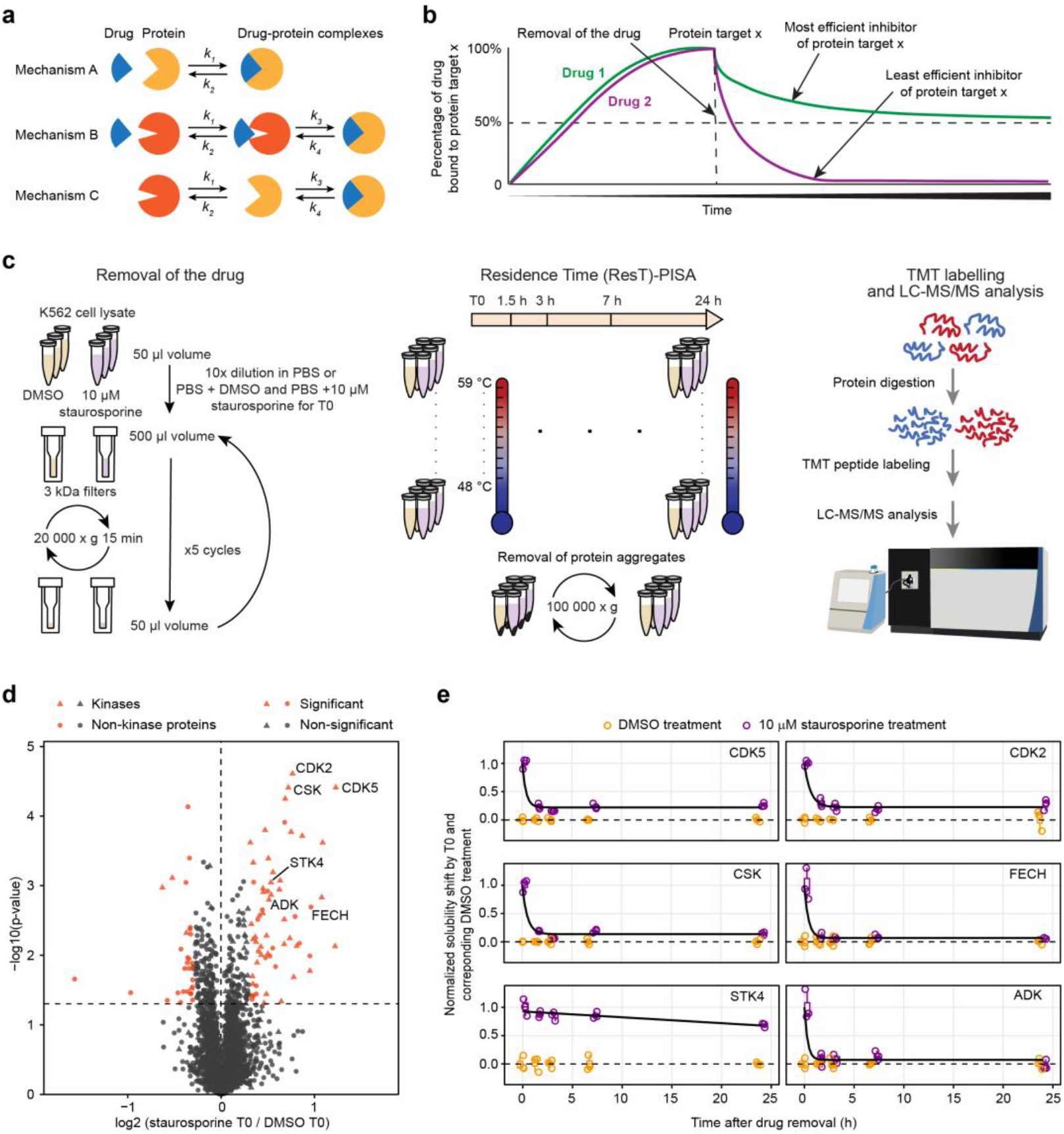
Drug-target residence time assessment using PISA assay. (**a**) The different modes of ligand-protein binding. (**b**) Residence time of drugs can vary significantly even when dose responses are similar for two different drugs (drug 1 and drug 2). (**c**) Workflow of ResT-PISA analysis using staurosporine treatment of K562 cell lysate. (**d**) Volcano plot of proteins solubility shift (ΔSm) in PISA between staurosporine and DMSO control at the start of the experiment (T0) corresponding to the maximum drug concentration. (**e**) Off-curves for selected targets show significant solubility changes with time past staurosporine removal. Horizonal line in the boxplots represent the median, 25th and 75th percentiles and whiskers represent measurements to the 5th and 95th percentiles. (n=3). *P*-values were calculated using a two-tailed Student’s *t*-test.

The above approaches are developed for studying simple systems with highly purified recombinant targets and drug molecules often modified for targeted purification or better detection. There exist proteome-wide methods of probing protein-target interactions not requiring drug modification, such as CETSA^18^, TPP^19^, as well as DARTS^20^ and LiP-MS^21,22^. These methods have been extensively used to deconvolute drug targets in cell lysate, intact cells, bodily fluids and even tissue. However, all these methods suffer from high analysis cost and low throughput. Some time ago we have introduced Proteome Integral Solubility Alteration (PISA) assay as a high-throughput analogue of TPP^23^. PISA analysis was used for deconvoluting the action mechanism of redox modulating anti-cancer compounds^24^ and profiling stem cell transitions^25^. Here we hypothesize that the ΔSm solubility shift in PISA will reflect the degree of target engagement by a drug in a cell lysate or intact cell at certain time past drug removal by filtration (for lysate) or washing (for cells). If so, the high throughput nature of PISA analysis should allow us to probe the residence time of a drug through the temporary profiles of ΔSm for different targets and off-targets within the same proteome. We also hypothesize that thus obtained residence times will be independent of the maximum ΔSm values that reflect the effect of drug binding on target solubility (sometimes interpreted, as thermal stability^19^). The mutual independence of these parameters would mean that addition of the residence time information to ΔSm data would augment the model predicting *in vivo* drug efficacy.

Here we test the above hypotheses by developing a novel technique called Residence Time Proteome Integral Solubility Alteration (ResT-PISA) assay. We show that most, but not all, proteins targets of staurosporine recover their solubility after a short period of time following drug removal, providing a proof of principle for ResT-PISA. Thereafter we verify that even covalent binding of a simple chemical group results in a complex residence time behavior reflecting both short-term binding as well as long-lasting interaction. To increase the throughput of our method, we compress ResT-PISA to cResT-PISA analysis and include dose response measurements (conc-PISA) in one multiplexed sample. This allowed us to create a scoring system based on three parameters - ΔSm, ResT and dose response, - to improve characterization and prioritization of efficacy targets. Lastly, we show that ResT-PISA is amenable to intact cells. Our new approach for evaluating drug target residence time in cell lysates and intact cells using an unbiased, label-free and system-wide method can find applications in drug discovery and development as well as in personalized medicine.

## Results

### ResT-PISA assay estimates drug-target residence times in a complex protein mixture

As a way of obtaining a proof of principle for ResT-PISA, we asked a question whether the multiple targets of a promiscuous drug will show different residence times in ResT-PISA analysis. To this end we treated a K562 cell lysate with 10 μM staurosporine chosen as the model drug known to inhibit many protein kinases^19^. We removed the drug after 45 min of treatment using 5 successive filtrations through 3 kDa membrane filters (approximately 1.10^6^ x dilution) (Fig. 1c). We opted for 3 kDa filters to minimize protein losses (Supplementary Fig. 1). Then we performed PISA analysis right after filtration (which corresponds to 1.5 h time interval from the start of the filtration process), as well as after 3 h, 7 h and 24 h. To obtain a measure of the effect of the maximal drug concentration on the solubility of protein targets, we analyzed by PISA a lysate treated with staurosporine and filtered with addition of PBS containing the same initial concentration of staurosporine (10 μM), in comparison with a control treated with vehicle (DMSO) instead of the drug (Fig. 1c). PISA measurements were also used to normalize the ResT-PISA results after removal of the drug. We recorded the off-curves for 5556 proteins quantified across all samples without missing values, among which 47 kinases showed significant solubility alteration after treatment with staurosporine at T0 (Fig. 1d). Most of these proteins recovered their solubility to the levels of an untreated sample after a short period of time past drug removal (Fig. 1e). These recovered proteins include multiple kinases, such as CDK2 and CDK5, and a common non-kinase off-target of kinase inhibitors, Ferrochelatase (FECH)^19,26^. However, some proteins like STK4 recovered less than half of their solubility change in 24 h after drug removal, suggesting a strong or covalent interaction (Fig. 1e).

### ResT-PISA analysis can detect binding events with prolonged lifetime

Being intrigued by the remarkably slow recovery of the staurosporine target STK4, we decided to investigate the off-curves of covalently attaching molecules. To this end we performed a ResT-PISA experiment in K562 cell lysate with 2 μM iodoacetamide (IAA) instead of staurosporine. IAA is widely used in proteomics to stabilize reduced cysteines by covalently adding to them a carbamidomethyl group. Since such alkylation reaction targets all accessible free cysteines, we expected to observe significant solubility shifts ΔSm for many proteins. We also expected that most of these proteins will show no or minimal recovery of solubility hours after removal of the chemical due to the covalent nature of the modification.

The results showed that the solubility was significantly affected by IAA in 165 proteins, of which 67 proteins lowered their solubility and 98 proteins (59%) increased it. While some of these proteins demonstrated, as expected, no solubility recovery 7 h after drug removal (Figs. 2a and b), some of the proteins did recover their initial solubility, suggesting that they either lost their cysteine modification or IAA was binding to them non-covalently (such binding is often the first step before covalent attachment). This observation highlights the complexity of protein-small molecule interactions that can happen through different modalities, as depicted in Fig. 1a. Interestingly, some of the targets of IAA with high ΔSm values and long residence times include two acetyl-COA acetyltransferases ACAT1 and ACAA2 (Figs. 2a and b). Since IAA has been used in molecular biology to inhibit glycolysis^27^, and as ACAT1 activity is linked to Warburg effect in cancer cells^28^, inhibition of acetyl-COA acetyltransferases may at least in part be responsible for IAA-induced reduction in glycolytic activity.

**Fig. 2.**
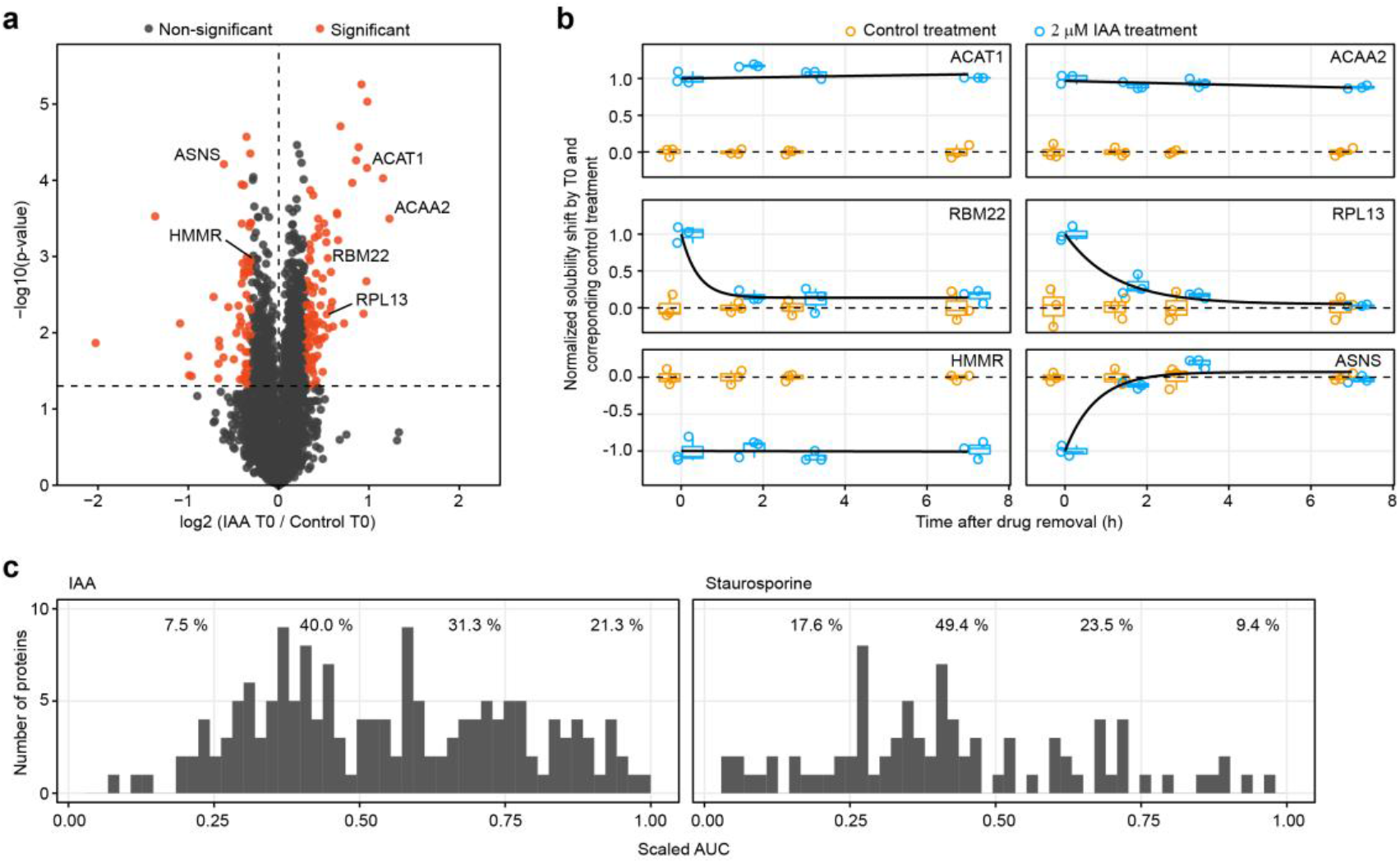
ResT-PISA of IAA as a covalent modifier. (**a**) Volcano plot of proteins’ solubility shift between IAA and water control at the start of the experiment (T0) corresponding to the maximum compound concentration. (**b**) Off-curves for selected targets show a significant solubility change with time past IAA removal. Horizonal line in the boxplots represent the median, 25th and 75th percentiles and whiskers represent measurements to the 5th and 95th percentiles. (**c**) Histograms of areas under the curves (AUCs) in (b) normalized by the maximum AUC value (n=3). *P*-values were calculated using a two-sided Student’s *t*-test.

To get further insight into the ResT-PISA specifics of the covalently attaching IAA and non-covalently interacting staurosporine, we compared for both drugs the distributions of the normalized areas under the curve (AUCs) that scale for a given drug with its residence time on a target protein. While AUCs of the majority (53%) of the detected targets of IAA scaled above 0.5, indicating slow-unbinding targets, for staurosporine the corresponding figure was significantly lower, 30% (Fig. 2c).

### Residence time is independent of the solubility shift ΔSm

Next, we tested ponatinib, a multi-target kinase inhibitor used in the clinic for treatment of chronic myeloid leukemia and Philadelphia chromosome-positive acute lymphoblastic leukemia^29^. PISA results revealed solubility shifts in multiple kinases previously reported as targets of ponatinib, such as LYN, MAPK14, CSK, IRAK4 and RIPK2^30^, as well as in kinases that have not yet been reported as ponatinib targets (Fig. 3a). Additionally, we detected a significant solubility shift for FECH. The residence time was assessed for all significantly shifting proteins as AUC of their off-curves (Fig. 3b). For all three molecules tested until this point, ΔSm did not correlate with the residence times of the proteins with shifting solubilities (Fig. 3c): the Pearson correlations of 0.02, -0.01 and 0.14 proved that these two parameters are largely independent. This result supported our hypothesis that residence time can be used as an independent kinetic parameter for target prioritization.

**Fig. 3.**
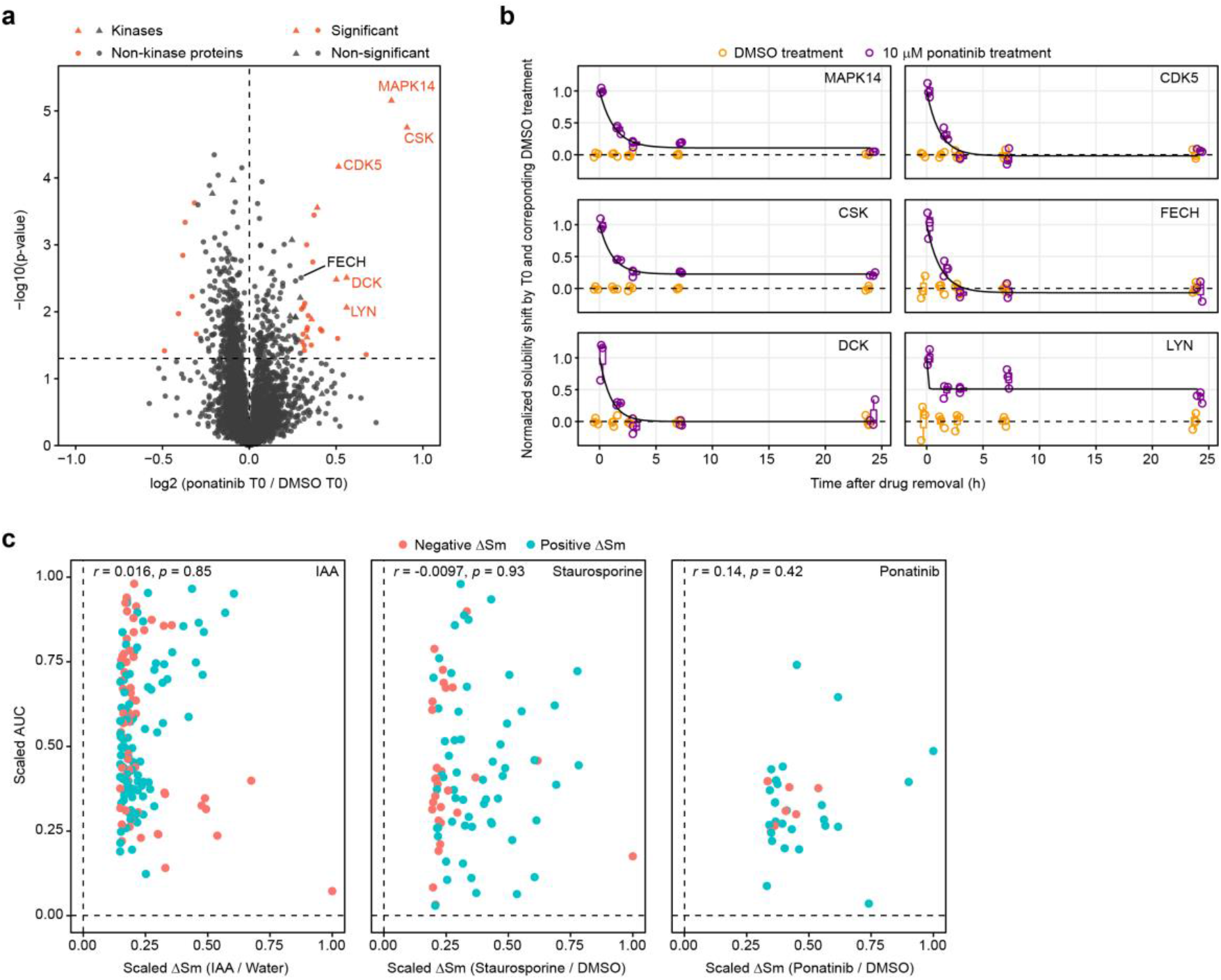
ResT-PISA of a kinase inhibitor ponatinib and comparison of AUCs with ΔSm. (**a)** Volcano plot of proteins solubility shift between ponatinib and DMSO control at the start of the experiment (T0) corresponding to the maximum drug concentration. (**b**) Off-curves for selected targets show a significant solubility change with time past ponatinib removal. Horizonal line in the boxplots represent the median, 25th and 75th percentiles and whiskers represent measurements to the 5th and 95th percentiles. (**c**) Scaled solubility shift ΔSm (x-axis) versus the residence time (y-axis) for proteins with shifting solubility in IAA, staurosporine and ponatinib treatments. Pearson correlation between the two measures were calculated for each treatment, n=3. *P*-values were calculated using a two-sided Student’s *t*-test.

### Off-curve measurements in ResT-PISA can be compressed

We investigated whether the ResT-PISA analysis could be further compressed by pooling all the time points together during sample preparation. This approach is similar to the one used in PISA in that it provides an AUC directly from one measurement. We called this approached compressed ResT-PISA or cResT-PISA. Since cResT-PISA can produce a triplicate analysis using only 9 samples (3 x PISA at T0, 3 x Control, 3 x cResT-PISA), we could multiplex two cResT-PISA analyses in a single TMT18-plexed set. The AUC from cResT-PISA correlated well (*r* =0.79) with the ResT-PISA AUCs for protein kinases (Supplementary Fig. 2), confirming that pooling the samples preserves residence time information.

Alternatively, we could multiplex with cResT-PISA altogether different measurements, such as the dependence of ΔSm upon drug concentration (Fig. 4a). Earlier we showed that this dose response curve can also be compressed in a PISA-like fashion^23^; here we compressed in “conc-PISA” 10 concentration points starting from 2 μM and proceeding with a 10-fold dilution at every successive concentration point. The results of the conc-PISA measurements provide a proxy for the association rate constant k_on_.

**Fig. 4.**
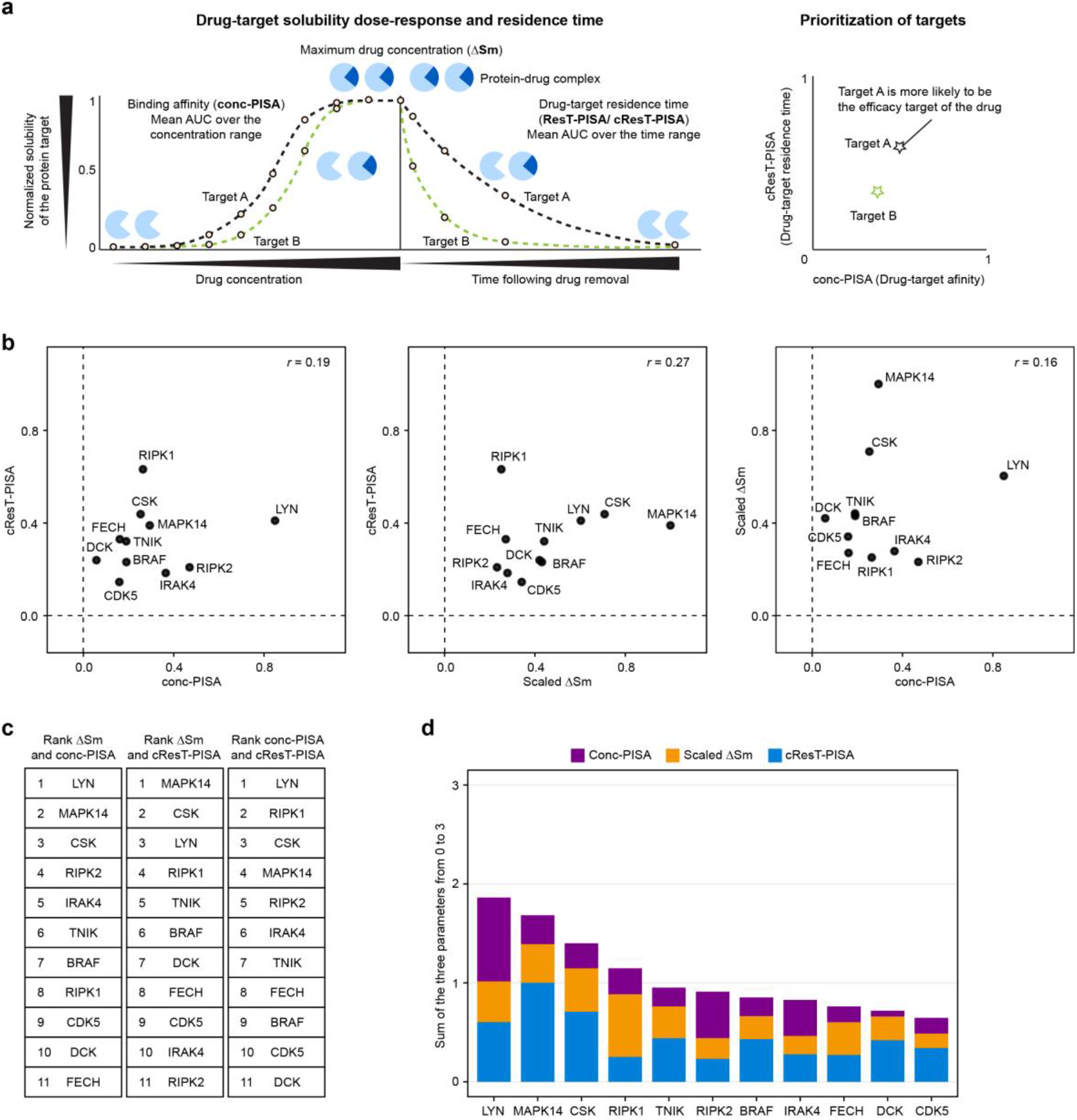
Combining PISA, cResT-PISA and conc-PISA results in a single score for prioritization of drug targets. (**a**) An example of comparing the dose responses and residence times and subsequent prioritization of two protein targets. (**b**) Plots of ΔSm against cResT-PISA, ΔSm against conc-PISA and cResT-PISA against conc-PISA for selected targets of ponatinib in K562 cell lysate. Pearson’s correlations were calculated for each plot. (**c**) Ranking of selected targets of ponatinib for ΔSm, cResT-PISA and conc-PISA scaled to the interval between 0 and 1. (**d**) Ranking of each target of ponatinib according to their combined scores.

We could multiplex in a TMT18 set three replicates of the following ponatinib analysis in K562 cell lysate:

1. Control PISA (incubation of lysate with DMSO for 45 min); no drug removal by filtering;
2. PISA of the merged samples incubated with ponatinib for 45 min in a range of concentrations; no drug removal by filtering;
3. PISA of the lysate incubated with ponatinib for 45 min at a maximum concentration of 2 μM; no drug removal by filtering;
4. Same but ponatinib is removed by spinning through a 3 kDa filter immediately upon incubation;
5. ResT-PISA control: same, incubated with DMSO instead of ponatinib;
6. PISA of the merged samples incubated with 2 μM ponatinib for 15 min, followed by drug removal by spinning through a 3 kDa filter with a varying time delay upon incubation.

### Combining PISA, cResT-PISA and conc-PISA results in a single score for target prioritizing

We compared the relative solubility shifts in PISA, cResT-PISA and conc-PISA and found no apparent correlation between them, confirming that these parameters are largely independent (Fig. 4b). Rankings of each target by the corresponding parameter are shown in (Fig. 4c). In order to combine the relative contribution of each parameter, we scaled each of them to a minimum of 0 and a maximum of 1 and summed the results to obtain a final score and overall target ranking. The known target of ponatinib ABL1 was not among the top ranked proteins, as we did not detect significant solubility shift in that protein. This could be due to the fact that K562 cells express a BCR-ABL fusion protein which earlier showed no ΔTm shift in TPP^19^. Instead, LYN kinase, a well-known target of ponatinib, ranked first. LYN is engaged by ponatinib at one of the lowest concentrations among known ponatinib targets^30^; in addition, it is known to produce a substantial thermal stability shift in TPP^30^. LYN was found by ResT-PISA to associate with ponatinib for a prolonged time, which explains why this protein came up as the top target in our analysis. Two other top proteins in overall ranking are also known targets MAPK14 and CSK that engage ponatinib at low concentrations and exhibit large thermal stability shifts in TPP^30^.

### Visualization tool for X-PISA analyses

In order to facilitate analysis of a combined set of PISA data (PISA, cResT-PISA and conc-PISA) we designed a data processing and visualization tool that is available on Github (https://github.com/RZlab/PISA-Analyser). The user guide to the interface is provided in Supplementary Fig. 3.

### ResT-PISA is amenable to intact cells

Unlike cell lysate, a living cell actively imports, metabolizes and exports drugs and their metabolites, and thus being able to monitor drug-target residence time in a cellular environment would be very valuable for predicting *in vivo* drug efficacy. Thus, we performed ResT-PISA in intact K562 cells treated with staurosporine and compared the outcome with the lysate results. Since removal of the drug from the growth media is a simple washing procedure, we could start recording residence time as early as 20 min past drug removal, i.e. much faster than with a lysate (Fig. 5a). In addition, the cell experiment did not require extensive filtering that may result in protein loss. On the other hand, the cell experiment required normalization of each PISA dataset by separately measured protein abundance, as in PISA-Express^25^, and thus demands multiplexing of a larger number of samples. In order to meet this increased demand, we metabolically labeled proteins in cells with light and heavy Stable Isotope Labeling by Amino acids in Cell culture (SILAC)^31^ reagents and additionally used TMT16-plex labeling of tryptic peptides. We performed PISA-Express analysis at each time point using an expression control at T0 as well as after 1.5 h, 3 h, 7 h and 24 h past drug removal. The obtained thermal shifts as well as residence times provided a detailed picture of drug effects in intact cells (Fig. 5a).

**Fig. 5.**
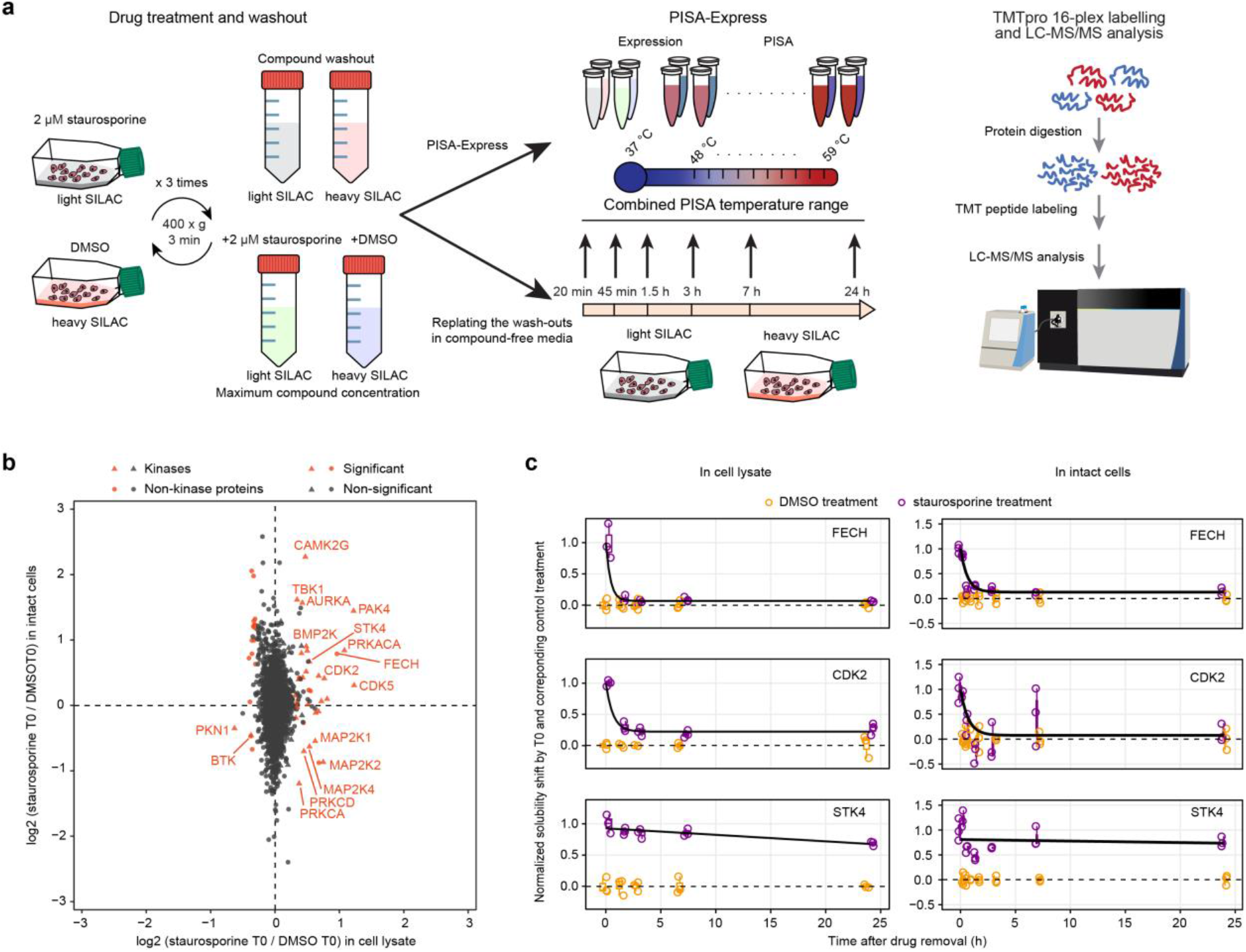
ResT-PISA in intact cells. (**a**) In-cell ResT-PISA workflow. (**b**) A comparison of ΔSm at T0 in K562 cell lysate (x-axis) and intact cells (y-axis) for staurosporine treatment against DMSO control. (**c**) Off-curves for three selected targets showing a significant solubility shift upon staurosporine removal in both cell lysate and intact cells. Horizonal line in the boxplots represent the median, 25th and 75th percentiles and whiskers represent measurements to the 5th and 95th percentiles. n=3. *P*-values were calculated using a two-sided Student’s *t*-test.

Surprisingly, some of the solubility shifts observed in lysate reversed their sign in cells (Fig. 5b); this was the case for MAP2K1, MAP2K2, MAP2K4, PRKCA, and PRKCD. MAP2Ks both phosphorylate kinases and are phosphorylated by kinases in signaling cascades involved in stress response^32^. PRKCA is a calcium activated and diacylglycerol-dependent protein, while PRKCD is phospholipid- and diacylglycerol-dependent; both proteins are involved in several regulatory processes including regulation of apoptosis^33^. These features may explain the difference in the direction of the solubility alteration in cells and lysate. When considering the absolute values of the solubility shifts, a positive correlation between the ResT-PISA AUCs in lysate and intact cells for common kinases was observed (r=0.59), while for all common proteins there was an anti-correlation (r=-0.39) (Supplementary Fig. 4). This result once again highlights the function of staurosporine as kinase inhibitor.

Many proteins showed similar residence times in cell lysate and intact cells (Fig. 5c). While for fast unbinding in lysate targets, such as FECH and CDK2, this result could be expected, it came as a surprise for slow unbinding targets, as the intact cell could detoxify the drug metabolically and/or degrade and replace the inhibited protein within hours past drug removal. However, STK4 which showed little recovery of solubility in lysate 24 h after removal of the drug had a similar behavior in intact cells, suggesting that STK4 stayed all this time bound with staurosporine and avoided being replaced in the cellular environment.

## Discussion

Here we developed ResT-PISA and its compressed version cResT-PISA, a compact approach to measuring drug target residence times at the proteome level in both cell lysate and intact cells. As well as other PISA-based techniques, ResT-PISA doesn’t require drug modification or the use of recombinant proteins. Importantly, the ResT-PISA approach is not limited to a selected class of compounds, as e.g. the competitive probes method that targets proteins with defined activity, such as kinases. Being multiplexed with PISA and concentration-dependent PISA analyses, this approach provides a comprehensive assessment of target engagement, binding affinity and residence time simultaneously. This enables extensive characterization of drug properties in a single analysis and provides a combined score for target prioritization. As a limitation, very short time frames (seconds or milliseconds) are not currently available due to the longer time required to remove the drug through cell washing or lysate filtration.

## Methods

### Cell culture

Chronic Myelogenous Leukemia K-562 cell line passage 3-8 (CCL-243™, ATCC) was maintained in non-adherent culture flasks (Sarstedt) in IMDM (Cat. No. L0190, Biowest) supplemented with 10% fetal bovine serum (Cat. No. 10500064, Gibco™).

#### Sample preparation for ResT-PISA in cell lysate

K-562 cells were grown until they reached a concentration of around 1.10^6^ cells/ml. Then 40 ml of cell culture were pelleted at 400 x g for 3 min and washed with PBS; the operation was repeated twice. Cells were pelleted one last time at 400 x g for 5 min and resuspended into 5 ml of PBS (GIBCO) supplemented with protease inhibitors (ThermoFischer Scientific). The cell lysate was clarified by centrifugation at 20 000 x g for 5 min and the supernatant was collected. 100 μl of supernatant were aliquoted in 30 PCR tubes, half were treated at a concentration of 10 μM of staurosporine (Selleckchem), 2 μM ponatinib (Selleckchem) or 2 μM iodoacetamide (IAA) (Sigma-Aldrich), the other half was treated with vehicle (DMSO or water) and kept at 37 °C for 45 min. 50 μl of each aliquot were transferred into a 500 μl 3 kDa filtration unit (Merck) and immediately 450 μl of PBS supplemented with protease inhibitors were added to each aliquot, with 3 aliquots receiving additionally the same initial concentration of a drug and 3 aliquots receiving the same concentration of vehicle instead. Then the filtering units were centrifuged at 17 000 x g for 15 min and 450 μl of the same solutions were added on top of each sample (10 x dilution). The operation was repeated 6 times. PISA assay was then performed on each sample at different time points, with T0 corresponding to the samples that were filtered with addition of a drug. Given the time required for centrifugation and aliquoting for PISA assay (see below), the first time point was 1.5 h after the start of drug removal. These samples were processed in parallel with the samples that received drug and DMSO during filtration (T0) for PISA analysis. The rest of the samples were left at RT for 3 h, 7 h and 24 h from the start of drug removal, and then were subjected to PISA analysis.

#### Sample preparation for ResT-PISA in cell

K-562 cells were grown in IMDM for SILAC (ThermoFischer Scientific) supplemented with either heavy (Cambridge Isotope Laboratories) or light (Sigma Aldrich) lysine and arginine and 10 % dialyzed FBS (ThermoFischer Scientific). Cells were grown until they reached a concentration of around 1.10^6^ cells/ml. Cells were split into two groups in 75 cm^2^ culture flasks (Sarstedt), the first group corresponding to heavy SILAC was treated for 1.5 h using 2 μM staurosporine and the other group corresponding to light SILAC was treated with the equivalent concentration of vehicle (DMSO) (Fig. 5a). For T0, triplicates of staurosporine or DMSO treatment were deposited in individual tubes and rinsed three times with PBS containing protease inhibitors as well as the corresponding concentration of staurosporine or DMSO, to control for potential changes happening in the samples during the centrifugation steps. For the samples corresponding to the 20 min time point, the cells were deposited in tubes in triplicates and rinsed three times with PBS containing protease inhibitors. Both cells from T0 and 20 min time point were then distributed into PCR tubes for PISA assay (see below). For longer time points, cells treated with staurosporine or DMSO were rinsed three times with fresh medium. Then the cells were seeded into 25 cm^2^ culture flasks (Sarstedt) in triplicates for each treatment and time point and placed again in the cell incubator for the intended duration. Then the cells previously treated with staurosporine or DMSO were rinsed three times with PBS, aliquoted into PCR tubes and processed for PISA assay (see below).

#### Compressed ResT-PISA combined with dose-response PISA (conc-PISA)

For cResT-PISA, ResT-PISA analysis in K562 cell lysate was performed as described above with the only difference being that after heating the aliquots and snap-freezing, the temperature and time points corresponding to one replicate of ResT-PISA were all pooled together, which resulted in 9 samples: 3 × 2 μM ponatinib treatment, 3 x DMSO treatment, 3 x of the pooled ResT-PISA range (cResT-PISA). In parallel, we performed conc-PISA analysis as previously described^23^. Briefly, the same K562 cell lysate was aliquoted into 12 samples per replicate and treated with the following concentrations for 45 min: 2 samples with 2 μM, 2 samples with DMSO only, one sample each with 2 pM, 20 pM, 200 pM, 2 nM, 20 nM, and 200 nM. Then each sample was aliquoted into of 9 tubes in triplicate and each replicate was heated for 3 min to the same 9 temperature points as in ResT-PISA in then left at RT for 3 min and snap-frozen. After this all 9 samples in each replicate were pooled together into a conc-PISA replicate. For the maximal and minimal drug concentration, only the samples corresponding to the treatment with 2 μM of ponatinib and DMSO were pooled together. This resulted in 9 samples in total: 3 × 2 μM ponatinib treatment, 3 x DMSO treatment, 3 x of the pooled concentration range (conc-PISA). After pooling the samples were processed as in PISA analysis (centrifugated and digested for LC-MS/MS analysis).

### PISA assay

Each condition was analysed in triplicate. For each replicate, 20 μl of cell suspension were aliquoted into 9 PCR tubes and each was heated for 3 min at a different temperature ranging from 48 °C to 59 °C. The samples were then kept for 3 min at RT before snap freezing in liquid nitrogen. After that, aliquots corresponding to each temperature point were combined. For ResT-PISA in cell, one sample designated for protein expression measurement was incubated at 37 °C (n=3 for each condition) and processed alongside the pooled samples. Cells were lysed using repeated freeze/thaw cycles and aliquots were combined as for the lysate experiment. Finally, all samples were transferred to ultracentrifuge tubes, placed into a Ti 42.2 rotor (Beckman-Coulter), and centrifuged at 100 000 x g for 20 min using an Optima XPN-80 Ultracentrifuge (Beckman-Coulter). 70 μl of the supernatant were collected and the same volume of lysis buffer (8 M urea, 20 mM EPPS pH 8.5) was added. The protein concentration was measured using Pierce bicinchoninic acid assay (BCA) protein assay kit (Thermo Fischer Scientific) according to the manufacturer’s protocol. The volume of sample was adjusted to the one corresponding to 25 μg of protein and the samples were processed for MS analysis (see below).

#### Protein sample preparation for expression proteomics and TMT labeling

For all proteomics experiments, 20 μg of proteins were used in sample preparation. S-S bond reduction was performed using 5 mM DTT at RT for 1 h followed by alkylation using 15 mM IAA at RT in the dark. The reaction was quenched by adding 10 mM of DTT. Then methanol/chloroform precipitation was performed as follows: 3 sample volume of methanol were added, then 1 sample volume of chloroform and 3 volumes of water. Samples were vortexed between each step and then centrifuged at 20 000 x g for 10 min at 4 °C. The aqueous layer was removed, and the protein pellet was rinsed with one sample volume of methanol, vortexed and centrifuged using the same speed as in the previous step. Finally, all the liquid was removed, and the protein pellet was air-dried.

Air-dried protein pellets were resuspended in 8 M urea, 20 mM EPPS pH 8.5. The samples were diluted once by adding 20 mM EPPS pH 8.5 (4 M urea), and lysyl endopeptidase digestion was carried out at a 1:100 ratio (LysC/protein, w/w) overnight at RT. The following day, samples were diluted 4 times (1 M urea) with 20 mM EPPS pH 8.5, then tryptic digestion was performed for 6 h at RT using a 1:100 ratio (Trypsin/protein, w/w). For PISA analysis, a “linker” corresponding to a sample composed of one tenth of each sample pooled together was prepared for normalization purpose. After that, TMT11, TMT16 or TMT18 labeling was performed for 2 h at RT by adding 0.2 mg of reagent dissolved in dry ACN according to manufacturer’s instructions. The ACN content in the samples was adjusted to a final concentration of 20%. The reaction was then quenched by adding triethylamine to a final 0.5% concentration. The samples were incubated for 15 min at RT and all temperature points were combined into one pooled sample per replicate. The pooled samples were acidified to pH < 3 using TFA, desalted using Sep Pack (Waters) and vacuum dried overnight using miVac DNA (Genevac).

#### High pH reversed-phase peptide fractionation

150 μg of peptides were resuspended in 20 mM NH_4_OH. Then, samples were off-line high-pH reversed-phase fractionated as described previously^34,35^ using an UltimateTM 3000 RSLCnano System (Dionex) equipped with a XBridge Peptide BEH 25 cm column of 2.1 mm internal diameter, packed with 3.5 μm C18 beads having 300 Å pores (Waters). The mobile phase consisted of buffer A (20 mM NH4OH) and buffer B (100% ACN). The gradient started from 1% B to 23.5% in 42 min, then moved to 54% B in 9 min, 63% B in 2 min and stayed at 63% B for 5 min, and finally moved back to 1% B and stayed at 1% B for 7 min. The fractions were collected for 0.7 min each resulting in 96 fractions that were concatenated into 24 fractions (fraction 1 was pooled with fractions 25, 49 and 73, fraction 2 - with fractions 26, 50 and 74 and so on) and dried using miVac DNA (GeneVac, England).

#### Mass spectrometry analysis

Each sample was resuspended in 2% ACN and 0.1% FA at a concentration of 0.2 μg/μl and 1 μg was injected into the respective LC system (summarized in Supplementary Table 1). Mass spectra were acquired using the parameters listed in Supplementary Table 1.

### Quantification and Statistical Analysis

#### Mass spectrometry data analysis

Raw files were converted to mzML format by MSConvert (version 3.0.21258)^36^. Peak picking of profile spectra was performed with the vendor-provided algorithm (Thermo Fisher Scientific). Then individual datasets were searched using FragPipe GUI v17.1 with MSFragger (version 3.4)^37^ as the search algorithm. Protein identification was performed with the human Swisspot database (20‵409 entries, downloaded on 2022.02.22), with acetylation (N-terminus) and oxidation on methionine as variable modification and carbamidomethylation of cysteine residues, TMT or TMTpro on the N-terminus or lysine as fixed isobaric labels. Trypsin was set as the enzyme with up to two missed cleavages. The peptide length was set to 7 – 50, and peptide mass range to 200 – 5000 Da. For MS2-based experiments, the precursor tolerance was set to 20 ppm and fragment tolerance to 20 ppm. Peptide-spectrum matches (PSMs) were adjusted to a 1% false discovery rate with Percolator^38^ as part of the Philosopher toolkit (v4.1.0+)^39^. For TMT labeled samples, reporter intensities were extracted by TMT-Integrator with default settings. As TMTpro18-plex labeling was not supported, reporter intensities were extracted by a home-written algorithm in R (version 4.1.1) as follows: 20 ppm windows around the theoretical m/z of each reporter ion were investigated, and the abundance of the most intense peak was extracted. For the SILAC-TMT all settings were the same, expect for the fragment tolerance that was set to 0.6 Th; also, heavy Lysine (+8.014199 Da) and Arginine (+10.008269 Da) were added as variable modifications.

#### Bioinformatics analysis

All further data processing was performed by a home-written algorithm in R (version 4.1.1). For MS2-based experiments, protein quantification was performed as follows: 1) PSMs mapping to a reverse sequence, known contaminants, with a precursor purity below 0.5 or a PeptideProphet probability below 0.9 were removed. 2) Reporter ion intensities were adjusted to correct for the isotopic impurities of the different TMTpro reagents by solving a system of linear equations according to manufacturer’s instructions. PSMs with corrected TMT intensities below 1000 were removed. 3) If multiple PSMs were detected in the same fraction with the same charge state, the one with the highest purity was retained, while for PSMs with the same purity the one with the highest Hyperscore was selected. 4) Individual protein TMT reporter intensities were calculated as the sum of the individual PSMs. 5) Proteins with less than two unique peptides were removed. 6) To correct for pipetting error, protein TMT reporter intensities were normalized by median centering to the total intensity. 7) Batch effects between different TMT sets were corrected by dividing each protein’s TMT intensity by their corresponding linker value. For the MS3-based SILAC-TMT experiments, protein intensities were calculated the same way as for the MS2 experiments; only PSMs having both the heavy and light version were quantified.

#### Time course ResT-PISA

Protein abundance at each time point was normalized by the corresponding vehicle value and statistical analysis was performed with a two-tailed t-test with equal variance. For visualization purposes, only proteins with a log2-scaled abundance fold change of at least 0.3 and a p-value below 0.05 were considered to be altered and used for further analysis of the residence time.

For curve fitting of residence time data, protein fold-changes were first scaled to the mean fold change of the drug-treated sample at time 0. Then, if the mean of the drug-treated sample at the last time point was below 0.6, an exponential off-curve with an asymptotic term:

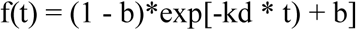

was fitted. If not, then a linear function

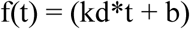

was fitted, where b – the asymptotic term, and k_d_ is the rate constant for protein-drug binding.

Area under the curve (AUC) calculations were made by summing for each replicate the mean of all data points over the entire time course.

Full curve ResT-PISA data in intact cells were treated the same way with the exception that changes in the protein expression were accounted for by normalizing each time point by their corresponding expression value.

All correlations were calculated as Pearson’s linear correlations.

#### Compressed concentration and residence time PISA

In both experiments, the protein abundance changes were normalized by those of the corresponding vehicle-treated samples. The significance of abundance changes was calculated as above. For further analysis the significant proteins from the maximal drug concentration or T0, for conc-PISA and cResT-PISA, respectively, were selected for analysis and protein solubility alterations were first scaled to the mean of their respective PISA samples treated with either ponatinib or DMSO. Then, proteins with a standard deviation between the replicates at the maximal concentration or first-time point bigger than three times the maximal mean value were removed. Lastly, proteins that had a scaled conc-PISA or ResT-PISA value above 1 or below 0 were removed.

The combined score for the filtered proteins was calculated as follows: 1) the scaled protein solubility alteration was determined by scaling each protein mean solubility alteration at the maximal drug concentration by the biggest absolute value among these proteins (alteration.score), 2) the scaled concentration value was calculated as described in 1) (concentration.score) and 3) the scaled residence time value was calculated as described in 1) (residencetime.score).

The final score was calculated as the sum of the scores in 1), 2) and 3).

## Data availability

The mass spectrometry proteomics data have been deposited to ProteomeXchange Consortium (http://proteomecentral.proteomexchange.org) via the PRIDE partner repository with data set identifier PXD034351.

## Code availability

The code for the web-interface is available on GitHub/RZlab (https://github.com/RZlab/PISA-Analyser).

## Supplementary figures

**Supplementary Fig. 1.**
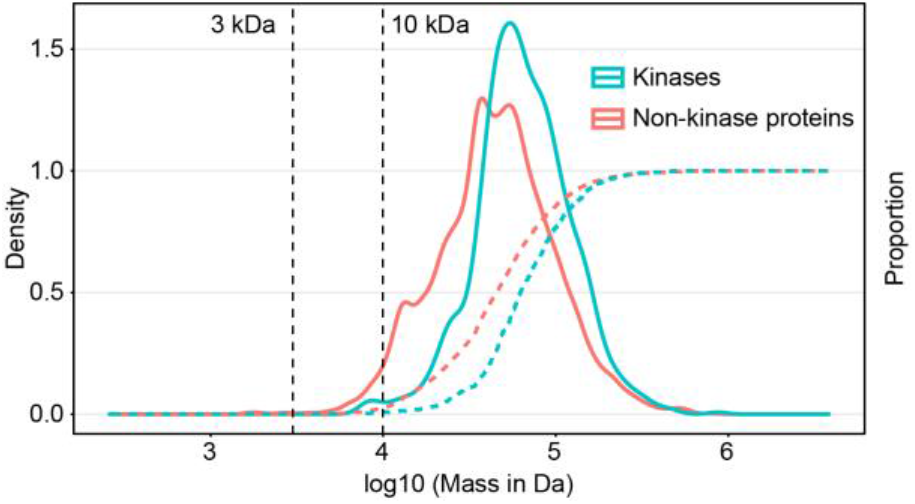
Protein mass distribution. Distribution of the molecular mass of protein kinases and other proteins (solid lines) and their proportion (dashed lines). The 3 kDa cutoff used for filtration for drug removal in this study is highlighted as well as another popular 10 kDa cutoff.

**Supplementary Fig. 2.**
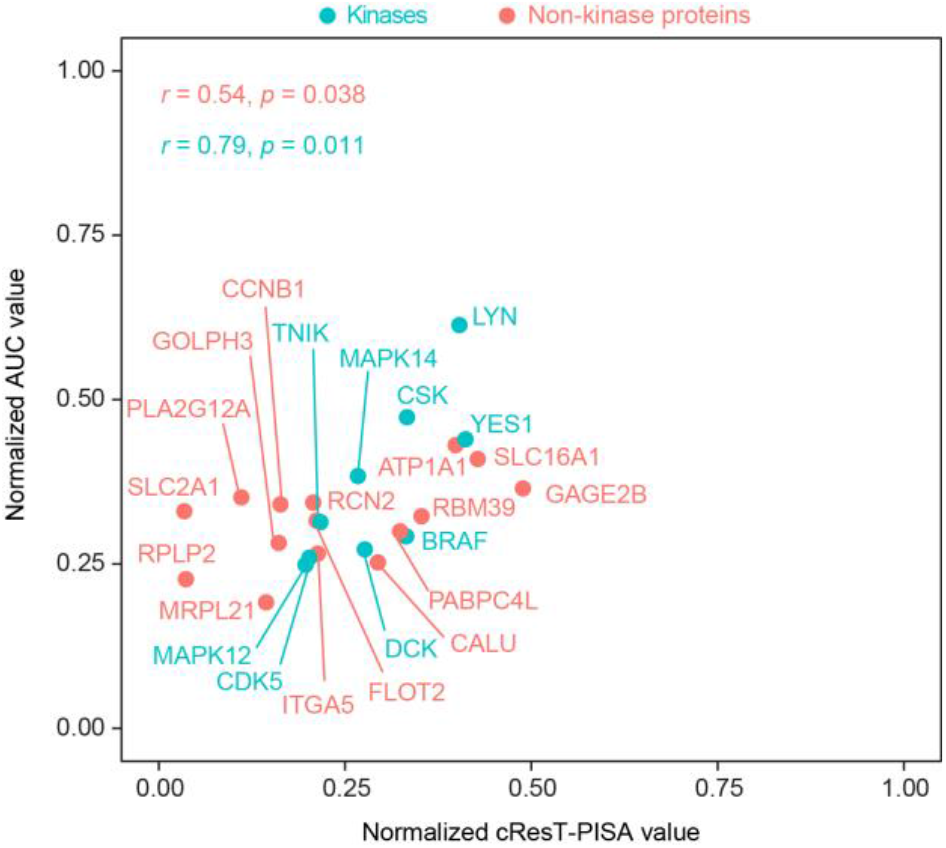
Correlation between the theoretical compressed ResT-PISA of ponatinib targets versus the observed experimental values. Protein kinases are shown in cyan and non-kinase proteins are shown in red. Pearson’s correlations between the two measures were calculated for the two protein groups.

**Supplementary Fig. 3.**
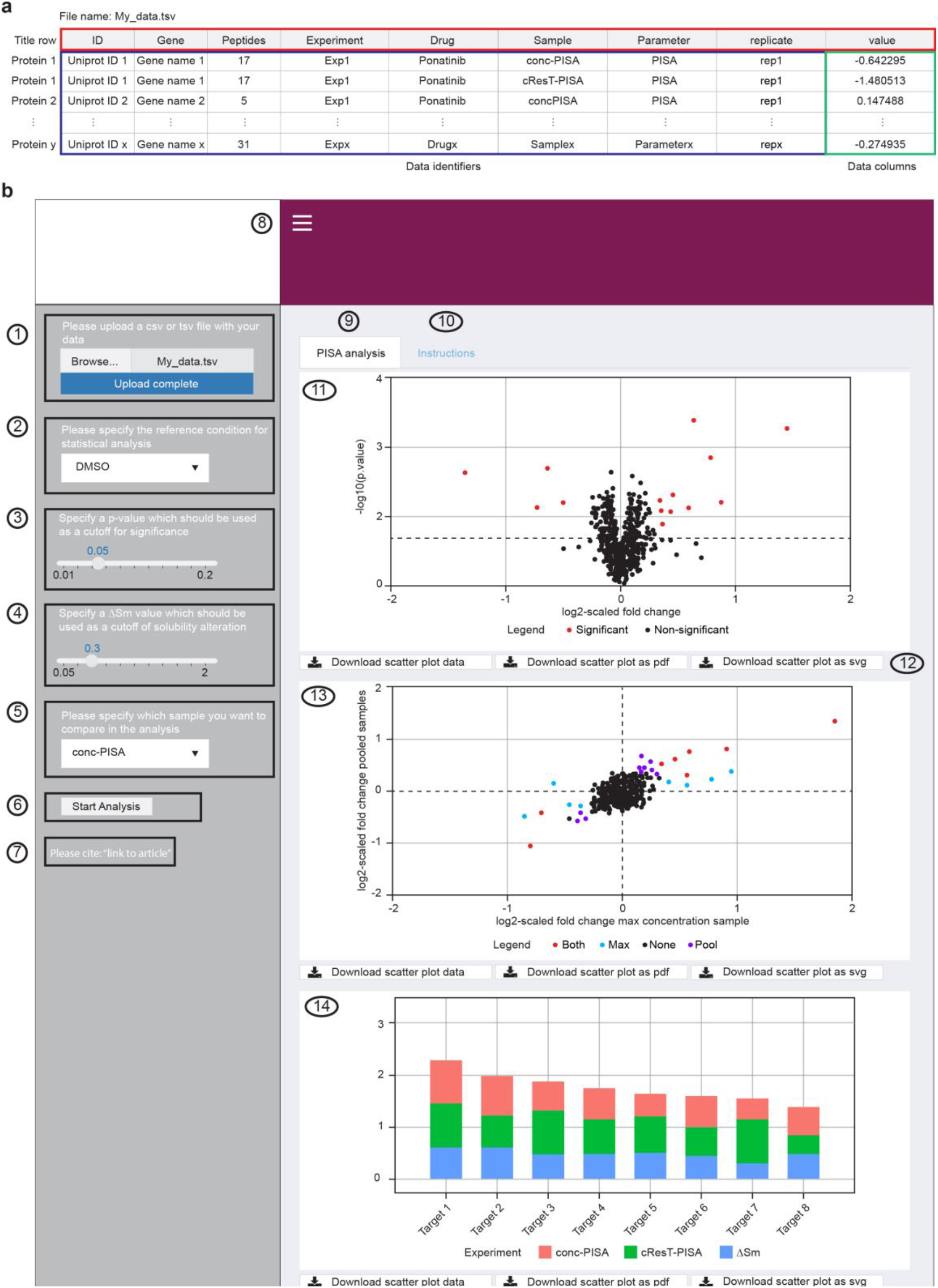
Overview of the PISA-analyzer interface. (**a**) Layout of a typical .tsv file containing a proteomics dataset that can be submitted to the interface for analysis. Data columns names should be presented as described here. Any number of conditions and replicates can be included in the .tsv file. Any condition name and replicate name can be included. (**b**) Step by step description of the web interface. 1) The user can upload own data here. 2) The user needs to specify the reference condition, in most cases the vehicle (DMSO here). 3) The user can specify the p-value cutoff for significance in the analysis. 4) The user can specify the ΔSm cutoff for significance in the analysis. 5) The user needs to specify which sample is to be compared to the reference sample in the analysis. 6) Start button of the analysis. 7) Link to the article. 8) Hide or show the checkbox group. 9) PISA analysis window where the user can see the result of his analysis. 10) Instruction window which displays the figure and caption. 11) Volcano plot of the sample chosen for comparison against the reference condition (from steps 2 and 5) presented as the mean log2 fold change (FC) (x-axis) and -log10 p-value (y-axis). The significant proteins according to the chosen p-value and ΔSm cutoffs are highlighted in red. 12) The plots can be exported as .pdf and .svg, the source data of the plots can also be downloaded as .tsv. 13) Optional plot that will appear only in 2-dimensional experiments (ResT-PISA or conc-PISA). Here two-dimensional plots of target engagement (in this case from maximum concentration of the drug ΔSm) and residence time (in this case from cResT-PISA analysis) presented as the mean log2 FC based on the chosen reference and sample in steps 2 and 5. 14) Optional plot that will appear only in 2-dimensional experiments. Here scoring based on the three dimensions of target engagement (maximum drug concentration), residence time (c-ResT-PISA) and binding affinity (conc-PISA). The score is calculated and is scaled to 1 for each individual parameter as described in the M

**Supplementary Fig. 4.**
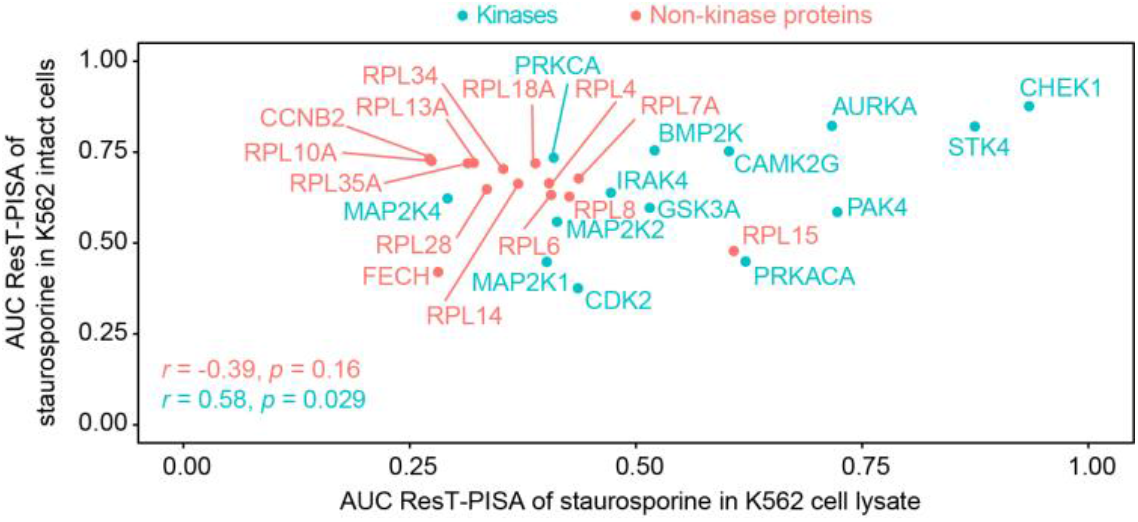
Correlation of ResT-PISA of staurosporine in lysate and in cell. Correlation between the AUC obtained from ResT-PISA of staurosporine in lysate and in cell. Kinase proteins are shown in cyan and non-kinase proteins in red. Pearson’s correlations between the two measures were calculated for the two protein groups.

**Supplementary Table 1.** LC-MS/MS systems and parameters.

| Parameter | Staurosporine Lysate | IAA Lysate | Ponatinib Lysate | Ponatinib PotPot | Staurosporine Cell |
| --- | --- | --- | --- | --- | --- |
| **LC system** | Ultimate 3000 | Ultimate 3000 | Ultimate 3000 | Ultimate 3000 | Ultimate 3000 |
| **Gradient (min)** | 110 | 210 | 180 | 180 | 135 |
| **Sample multiplexing strategy** | TMT11-plex | TMT10-plex | TMTpro16-plex | TMTpro18-plex | TMTpro16-plex +SILAC |
| **Mass Spectrometer** | Fusion Lumos | Fusion Lumos | Fusion Lumos | Fusion Lumos | Fusion Lumos |
| **Scan cycle (s)** | 3 | 3 | 3 | 3 | 2.5 |
| **MS1 resolution** | 120`000 | 120`000 | 120`000 | 120`000 | 120`000 |
| **MS1 scan range (Th)** | 400-1600 | 400-1600 | 375-1400 | 375-1400 | 375-1400 |
| **Injection time (ms)** | 50 | 50 | 50 | 50 | 50 |
| **MS1 AGC** | 4*10<sup>6</sup> | 4*10<sup>6</sup> | 1*10<sup>6</sup> | 1*10<sup>6</sup> | 1*10<sup>6</sup> |
| **Included charge states** | 2-6 | 2-6 | 2-5 | 2-5 | 2-5 |
| **Mass Difference** | NA | NA | NA | NA | 8.0142, 10.0083, 16.0284, 20.0165, 18.0225, 24.0426, 30.0248, 26.0367, 28.0307, Tolerance 10 ppm |
| **Exclusion duration (s)** | 60 | 60 | 60 | 60 | 45 |
| **MS2 Isolation windows (Th)** | 1.6 | 1.6 | 0.7 | 0.7 | 0.5 |
| **MS2 NCE** | 35 (HCD) | 35 (HCD) | 35 (HCD) | 35 (HCD) | 35 (CID) |
| **MS2 resolution** | 60`000 | 50`000 | 50`000 | 50`000 | Turbo (Ion Trap) |

|  | (Orbitrap) | (Orbitrap) | (Orbitrap) | (Orbitrap) |  |
| --- | --- | --- | --- | --- | --- |
| **Injection time (ms)** | 118 | 86 | 86 | 86 | 35 |
| **MS2 AGC** | 125`000 | 125`000 | 250`000 | 250`000 | 25`000 |
| **MS3 Isolation windows (Th)** | NA | NA | NA | NA | 1.3 |
| **MS3 NCE** | NA | NA | NA | NA | 45 (HCD) |
| **MS3 resolution** | NA | NA | NA | NA | 50`000 (Orbitrap) |
| **MS3 Injection time (ms)** | NA | NA | NA | NA | 86 |
| **MS3 AGC** | NA | NA | NA | NA | 250000 |

